# Fungal communities in sediments along a depth gradient in the Eastern Tropical Pacific

**DOI:** 10.1101/2020.06.26.173682

**Authors:** Keilor Rojas-Jimenez, Hans Peter Grossart, Erik Cordes, Jorge Cortés

**Author notes:** Correspondence: Keilor Rojas-Jimenez, Hans Peter Grossart.

## Abstract

Deep waters represent the largest biome on Earth and the largest ecosystem of Costa Rica. Fungi play a fundamental role in global biogeochemical cycling in marine sediments, yet, they remain little explored. We studied fungal diversity and community composition in several marine sediments from 16 locations sampled along a bathymetric gradient (from a depth of 380 to 3474 m) in two transects of about 1500 km length in the Eastern Tropical Pacific (ETP) of Costa Rica. Sequence analysis of the V7-V8 region of the 18S rRNA gene obtained from sediment cores revealed the presence of 787 fungal amplicon sequence variants (ASVs). On average, we detected a richness of 75 fungal ASVs per sample. Ascomycota represented the most abundant phylum with Saccharomycetes constituting the dominant class. Three ASVs accounted for ca. 63% of all fungal sequences: the yeast *Metschnikowia* (49.4%), *Rhizophydium* (6.9%), and *Cladosporium* (6.7%). Although we distinguished a cluster dominated by yeasts and a second cluster dominated by filamentous fungi, we were unable to detect a strong effect of depth, temperature, salinity, dissolved oxygen, and pH on the composition of fungal communities. We highlight the need to understand further the ecological role of fungi in deep-sea ecosystems.

## INTRODUCTION

Fungi existed in the oceans long before they conquered land, and within the oceans, the deep sea represents the largest biome on Earth. Therefore, it is crucial to study fungal diversity and ecology in deep-sea waters, for which there is a paucity of studies compared to the rest of the ocean. Detailed knowledge of deep-sea fungi is required to understand better the overall fungal contribution to marine food webs and biogeochemical cycles at the global scale (Manohar and Raghukumar, 2013; Barone et al., 2018; Drake and Ivarsson, 2018; Grossart et al., 2019; Román et al., 2019; Hassett et al., 2020).

Fungi are active members of deep-sea sediment communities (Pachiadaki et al., 2016; Morales et al., 2019), but in this ecosystem, they are far more poorly characterized than bacteria and archaea (Edgcomb et al., 2011; Nagano and Nagahama, 2012; Dekas et al., 2016; Xu et al., 2018). In deep waters, fungi are well adapted to the total absence of light, low temperatures, and high hydrostatic pressure. Fungal communities have been described in sediments of hydrothermal vents, methane-cold seeps, oxygen-minimum zones, and associated with other macro-organisms (Nagahama et al., 2011; Zhang et al., 2016; Batista-García et al., 2017). Furthermore, the subseafloor has been shown to represent a vast ecosystem where micro-aerobic respiration occurs and where large amounts of microbial life subsist, even hundreds of meters below the seafloor (D’Hondt, 2002; Roy et al., 2012; D’Hondt et al., 2015; Ivarsson et al., 2016a; Nagano et al., 2016).

Fungi in the deep-sea environment mainly survive on marine snow, which consists of organic matter derived from photosynthesis that takes place in the photic layer. In addition to performing aerobic respiration, fungi are capable of carrying out processes such as fermentation, sulfate reduction, methanogenesis (Lenhart et al., 2012; Orsi et al., 2013; Bochdansky et al., 2017), and possibly lithoautotrophy (López-García et al., 2003; Nealson et al., 2005; Ivarsson et al., 2016b). These metabolic processes may be more critical for fungi in deep waters since it has been observed that as depth increases, fungal populations exhibit a more multitrophic lifestyle (Li et al., 2019).

In recent years, there has been a growing interest in studying fungal communities in deep-sea environments using culture-dependent and, to an increasing extent, culture-independent methods. Abundant fungal populations have been observed in a variety of deep-sea locations such as asphalt seeps in Sao Paulo Plateau (Nagano et al., 2017), methane seeps in the Kuroshima Knoll (Takishita et al., 2006), hydrothermal vents in the Mid-Atlantic Ridge (Le Calvez et al., 2009; Xu et al., 2017), sediments of the Peru Trench (Edgcomb et al., 2011), the East Indian Ocean (Zhang et al., 2014), the High Arctic (Zhang et al., 2015), the Mariana Trench (Xu et al., 2016, 2018), the Yellow Sea (Li et al., 2016), the Mediterranean Sea (Barone et al., 2018), the Yap Trench (Li et al., 2019), and subsurface sediments in Suruga-Bay (Nagano et al., 2016).

Considering the enormous area to be explored for fungal diversity and function in deep-sea sediments, the existing studies are minimal and often lack an adequate spatial and temporal resolution (Grossart and Rojas-Jimenez, 2016; Grossart et al., 2019; Morales et al., 2019). Therefore, there is still a large number of geographical locations that have not yet been studied, including the Eastern Tropical Pacific (ETP). The deep-sea waters of the ETP constitute a particularly important ecosystem in Costa Rica since they represent about 90% of the whole territory (Cortés, 2016, 2019).

The Costa Rican ETP comprises a chain of mountains and submarine volcanoes across the subduction zone of the Cocos and Caribbean tectonic plates. Here, there is a high diversity of microhabitats (Lizano, 2001; Protti et al., 2012; Rojas and Alvarado, 2012) including methane seeps (Sahling et al., 2008; Levin et al., 2012, 2015). Previous studies have shown high endemism and diversity of macro- and microorganisms in this region (Rusch et al., 2007; Cortés, 2008, 2019; Rojas-Jiménez, 2018). Also, the Costa Rican ETP is part of a marine corridor that extends through Isla del Coco to the Galapagos Islands in Ecuador, which represents an essential site for the conservation and regeneration of marine species throughout the ETP (Cortés, 2012).

In this work, we have explored the diversity and composition of fungal communities in deep-sea sediments of the Costa Rican ETP. Two expeditions were carried out with transects of approximately 1500 km length each, and sediments were sampled at 16 locations at depths between 380 m and 3474 m. We extracted DNA from subsamples of each sediment core, sequenced the 18S rRNA gene of eukaryotes, and performed a subsequent bioinformatic analysis. This work confirms the high abundance and diversity of fungi in sediments of the ETP region. We expect that our results will support current efforts to conserve this region by providing a baseline of the high diversity of eukaryotic species and microhabitats found in its deep-sea waters.

## MATERIALS AND METHODS

We used Illumina sequencing of the eukaryotic 18S rRNA gene to characterize the fungal community composition in sediments along a depth gradient in the ETP of Costa Rica, across two transects of ca. 1500 km length each (**Figure 1**). All samples were collected with the permission of the Ministry of Environment and Energy of Costa Rica (SINAC-CUSBSE-PI-R-032-2018; R-070-2018-OT-CONAGEBIO). The RV *Atlantis* surveyed the Pacific continental margin of Costa Rica from October 24^th^ to November 5^th^, 2018, from the continental slope to the offshore seamounts across a subduction zone. In this region, several methane-rich seeps have been detected (Sahling et al., 2008; Levin et al., 2012, 2015). All sediment cores were collected by the human-occupied vehicle (HOV) *Alvin* equipped with mechanical, maneuverable arms. We analyzed eight sediment-cores from this expedition. The following year, the RV *Falkor* surveyed the seamounts extending from the mainland to the Isla del Coco National Park between January 6^th^-21^st^, 2019. This region comprises several seamounts and natural gas seeps and provides an important corridor for highly specialized biological communities occupying the area. The sediment cores were collected employing the remotely operated vehicle (ROV) *SuBastian*, which is also equipped with mechanical, maneuverable arms. We analyzed another eight sediment cores from this expedition. Further details of the sampling sites, dates, and environmental variables measured are shown in **Table 1**.

**Table 1.** Sites of the Eastern Tropical Pacific of Costa Rica sampled in this study, with the respective values of the environmental variables measured.

| Sample | RV | Site | Date | Depth [m] | Temp [°C] | Salinity [PSU] | DO [mg/L] | pH | Data sources* |
| --- | --- | --- | --- | --- | --- | --- | --- | --- | --- |
| A1 | Atlantis | Mound 12** | 24/10/18 | 996 | 5,06 | 34,57 | 1,10 | 7,62 | 1, 6 |
| A2 | Atlantis | Quepos Slide** | 25/10/18 | 380 | 11,75 | 34,76 | 0,20 | 7,71 | 1, 7 |
| A3 | Atlantis | Quepos Plateau | 26/10/18 | 2200 | 2,06 | 34,60 | 3,73 | 8,06 | 2, 4, 6 |
| A4 | Atlantis | Seamount 3 | 28/10/18 | 1383 | 3,35 | 34,60 | 1,67 | 7,70 | 2, 4, 6 |
| A5 | Atlantis | Mound 11** | 3/11/18 | 1024 | 4,83 | 34,57 | 1,24 | 7,67 | 6, 7 |
| A6 | Atlantis | Jaco Scar** | 4/11/18 | 1788 | 2,54 | 34,63 | 2,42 | 7,61 | 1, 6 |
| A7 | Atlantis | Parrita Seep** | 5/11/18 | 1410 | 3,41 | 34,60 | 2,21 | 7,71 | 6,00 |
| A8 | Atlantis | Quepos Plateau | 26/10/18 | 1873 | 3,50 | 34,61 | 3,11 | 8,06 | 1, 3 |
| F1 | Falkor | The Thumb** | 10/1/19 | 1072 | 4,54 | 34,58 | 1,22 | 7,69 | 4, 7 |
| F2 | Falkor | Parrita Scar | 11/1/19 | 1419 | 3,35 | 34,61 | 2,08 | 7,67 | 4, 5 |
| F3 | Falkor | Rio Bongo | 13/1/19 | 659 | 14,41 | 34,93 | 1,50 | 7,60 | 4, 7 |
| F4 | Falkor | Subduction Seep | 14/1/19 | 3474 | 1,88 | 34,66 | 4,20 | 7,71 | 4, 5 |
| F5 | Falkor | Seamount 5.5 | 15/1/19 | 1540 | 3,00 | 34,62 | 2,64 | 7,70 | 4, 5 |
| F6 | Falkor | Seamount 7 | 16/1/19 | 1320 | 4,11 | 34,59 | 1,80 | 7,67 | 4, 5 |
| F7 | Falkor | Coco Canyon | 18/1/19 | 950 | 5,02 | 34,57 | 1,40 | 8,12 | 4, 5 |
| F8 | Falkor | Mound Jaguar** | 25/1/19 | 1903 | 2,43 | 34,63 | 3,13 | 7,75 | 4, 5 |

\* 1. AUV Sentry sensors, 2. HOV Alvin sensors, 3. HOV Alvin Niskin bottle, 4. ROV SuBastian sensors, 5. ROV SuBastian Niskin bottle, 6. RV Atlantis CTD, 7. RV Falkor CTD
\*\* Seep areas

**Figure 1.**
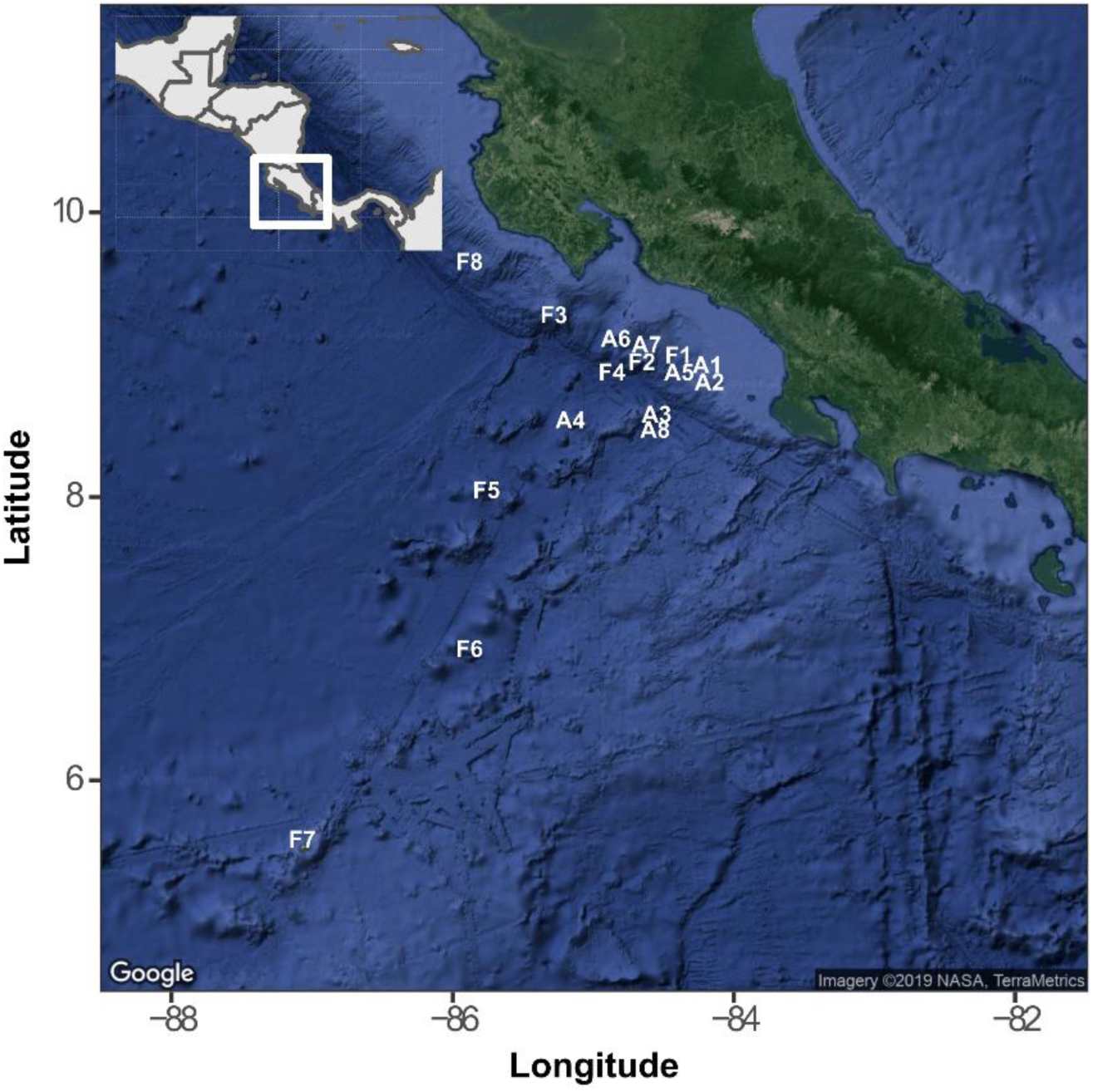
The geographical location of sampling points in the Eastern Tropical Pacific of Costa Rica. The points indicated with the letter **A** correspond to the route followed by the *Atlantis* cruise and those with the letter **F** to the *Falkor* cruise. The map was generated with the ggmap package using a Google satellite image.

After collection, nearly one gram of the upper (1-2 cm), middle (6-7 cm), and lower (13-14 cm) parts of each 15 cm-core was deposited into a 1.5 ml tube, stored at -20 °C on board the vessel and at -80 °C in the laboratory. The sediment DNA was extracted with a DNA isolation kit (PowerSoil^®^, Qiagen) following the manufacturer’s instructions. From some subsamples, unfortunately, it was not possible to obtain enough DNA for subsequent analyzes, so in total, we retrieved 40 DNA samples (out of the 48 possible) from the 16 cores sampled in both transects. For the construction of the amplicon library, primers FF390 / FR1 were used to amplify the V7 and V8 regions of the 18S rRNA gene (Prévost-Bouré et al., 2011). The products were subjected to a 250 nt paired-end sequencing using Illumina MiSeq technology at MrDNA, TX, USA.

Sequences were tested for quality and analyzed using version 1.12 of the DADA2 pipeline (Callahan et al., 2016). The taxonomic assignment was performed on comparing sequences against the SILVA reference database v132 (Quast et al., 2013), and then curated by comparing sequences against NCBI. Global singletons were removed. This process resulted in an amplicon sequence variant table, a higher-resolution analog of the traditional OTU table, which records the number of times each exact amplicon sequence variant was observed in a sample. Sequence data were deposited in the sequence-read archive (SRA) under accession PRJNA632873.

Statistical analyses and their visualization were performed with the R statistical program (R-Core-Team 2019) and the Rstudio interface. Package Vegan v2.5-6 (Oksanen et al., 2019) was used to calculate alpha diversity estimators, non-metric multidimensional scaling analyses (NMDS). Data tables with the amplicon sequence variants (ASV) abundances were normalized into relative abundances and then converted into a Bray–Curtis similarity matrix. To determine if there were significant differences between the fungal community composition according to factors such as depth or transect, we used the non-parametric multivariate analysis of variance (PERMANOVA) and pairwise PERMANOVA (adonis2 function with 999 permutations). For the network analysis, we selected the 24 most abundant eukaryotic ASVs (10 classified as fungi), which corresponded to nearly 70% of the total number of eukaryotic sequences. We considered a valid co-occurrence event if the Spearman’s correlation coefficient was >0.5 (Junker, 2008). The network inference and visualization were performed with package igraph v1.2.4.2 using a Fruchterman-Reingold layout (Csardi and Nepusz, 2006).

Environmental data were compiled from the measurements obtained from analyses of the water samples or sensors nearest to the sediment cores as possible. These data come from a variety of sources. Temperature and salinity data were obtained from the conductivity-temperature-depth (CTD) sensors on the HOV *Alvin* and ROV *SuBastian*, which were also equipped with niskin bottles for water sampling. There was a dissolved oxygen (DO) optode on the ROV *SuBastian* as well as the autonomous underwater vehicle (AUV) *Sentry* which was deployed over some of the sites during the 2018 *Atlantis* expedition. Niskin rosettes with attached CTDs were also deployed from the *Atlantis* and *Falkor* over the sites, and the *Falkor* CTD had a DO optode as well. DO data were compiled from a combination of these sources. DO data for the samples from the 2018 *Alvin* dives were derived from either the *Sentry* data (if available from the site) or calculated from a curve fitted to the DO data obtained from the ROV *SuBastian* and *Falkor* CTD DO-depth profiles. DO data for the 2019 SuBastian push core samples was deterimined from SuBastian optode. The pH data were exclusively from the water samples obtained by the rosette deployed from the ship or the niskin bottles on the submersibles. Water samples were brought to room temperature and the pHT (total scale) was measured using an Orion 5 Star pH meter and glass electrode (ROSS Ultra pH/ATC Triode 8107BNUMD) in triplicate within 4 h of collection (Dickson et al., 2007).

## RESULTS AND DISCUSSION

Fungi constituted the most abundant group of eukaryotic organisms in the sediments of the ETP of Costa Rica, according to the analysis of the sequences of the 18S rRNA gene. We determined the presence of 787 fungal ASVs out of a total of 2707 eukaryotic ASVs. Fungi represented 59.72% of the 2,746,436 sequences analyzed (**Figure 2A**). Other abundant eukaryotic groups comprised Cercozoa and Ichthyosporea, which represented 24,1% and 6,75% of all eukaryotic sequences, respectively. The genus *Gymnophrys* was the most abundant within Cecrozoa, while an ASV related to *Pirum* was the most abundant within Ichthyosporea. It was not possible to provide a classification at the taxonomic level of the Kingdom for 2.63% of the sequences, which, however, contain 24.60% of all observed ASVs. This indicates the presence of a large number of rare species that have not yet been registered in reference databases, which suggests a high hidden eukaryotic diversity in the studied deep-sea sediments.

**Figure 2.**
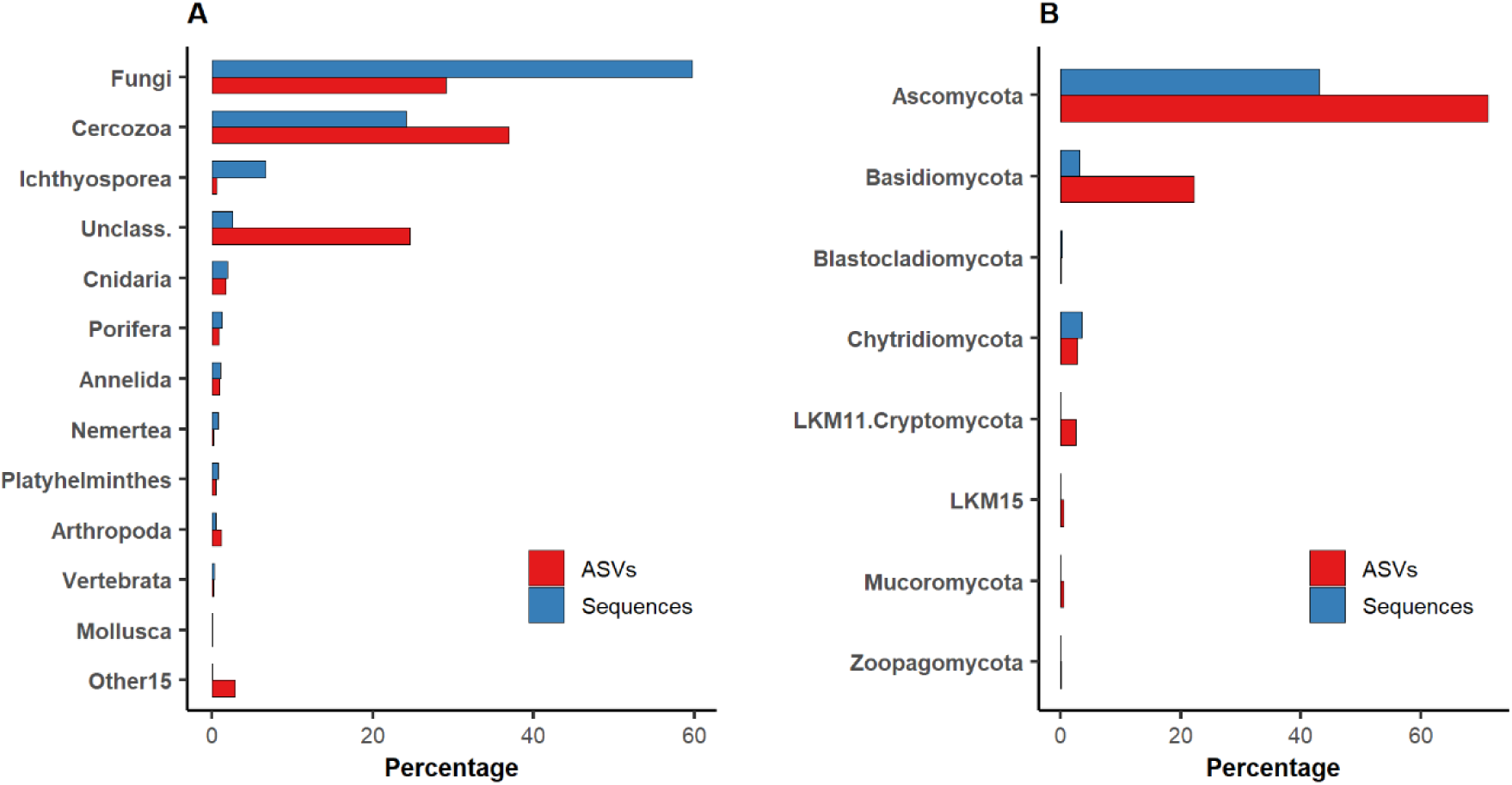
The relative abundance of eukaryotic groups (**A**) and fungal groups (**B**) in deep-sea sediments of the Eastern Tropical Pacific of Costa Rica concerning the number of sequences and amplicon sequence variants (ASVs).

The most abundant fungal phylum in marine sediments of the ETP of Costa Rica consisted of Ascomycota, which represented 43% of all fungal sequences and 71% of the ASVs. The second most abundant fungal group comprised Basidiomycota, representing nearly 3% of the sequences but 22% of the ASVs, suggesting a very high phylogenetic diversity within this phylum. Chytridiomycota represented the third most abundant fungal group, with 3.5% of the sequences and 2.79% of the ASVs. Other less frequent fungal groups observed in this ecosystem were, Blastocladiomycota, LKM11, LKM15, Mucoromycota, and Zoopagomycota (**Figure 2B**). These results are consistent with earlier results obtained in deep-sea sediments from several parts of the planet confirming the general dominance of Ascomycota in deep-sea sediments together with the presence of Basidiomycota and Chytridiomycota in lower proportions (Li et al., 2016, 2019; Xu et al., 2016, 2019; Zhang et al., 2016; Nagano et al., 2017; Barone et al., 2018; Wang et al., 2019).

When analyzing the relative abundances at the class level, we detected a total of 32 classes in the deep-sea sediments, where Saccharomycetes was the most prominent in the majority of the samples. In samples where Saccharomycetes was dominant, they were typically accompanied by the presence of Chytridiomycetes. There were also groups of samples with high abundances of Eurotiomycetes, Dothideomycetes, and Agaricomycetes, but where Saccharomycetes were practically absent (**Figure 3**).

**Figure 3.**
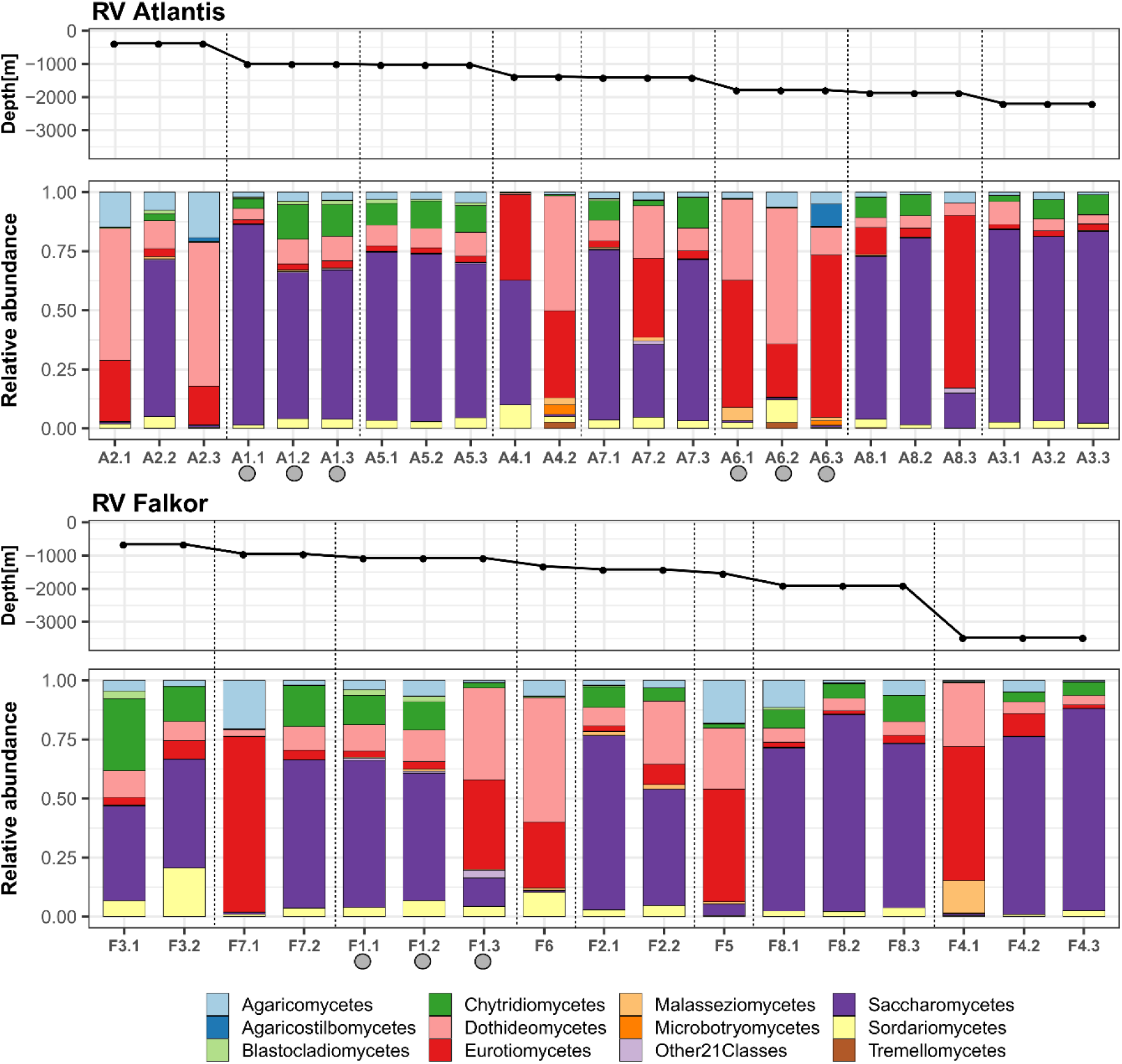
The relative abundance of fungi, at the taxonomic level of class, in deep-sea sediments of the eastern tropical Pacific of Costa Rica. The proportions within sampling points of the core subsamples for each of the cruise transects are shown. The samples were ordered according to the depth gradient. Gray circles indicate active methane seeps.

We also observed high variability in the composition within the horizons of some samples. In this regard, the homogeneity or heterogeneity of horizons could be related to the characteristics of the sampled habitat, but also with the sedimentation time. It will be necessary to further explore in more detail the variations that occur at the micro-scale in fungal communities in sediment profiles, as has been done to study variations in the physicochemical conditions of sediments in other deep-sea waters (Roy et al., 2012; D’Hondt et al., 2015; Román et al., 2019).

The samples of the deep-sea environment studied, characterized by high hydrostatic pressure, low temperatures, and the absence of light, presented an average richness of 75 fungal ASVs per sample (range 13-147), while the average value of the Shannon index was 1.77 (range 0.84-2.68). As with the community analyses, there were no significant differences in the alpha diversity estimations between depths and expeditions (Kruskal-Wallis, P> 0.05). The average value of the Pielou’s evenness was 0.42 (range 0.21-0.71), indicating a certain uniformity in the abundances of most of the observed phylotypes (**Supplementary Figure 1).**

The genus *Metschikowia* was the most abundant within the class Saccharomycetes and also the most abundant in the majority of the sediments analyzed. The genus *Metschikowia* comprises single-celled budding yeasts known for its participation in fermentation processes and wine production, reported mainly in terrestrial environments (Kang et al., 2017; Wang et al., 2017; Pawlikowska et al., 2019). In this study, we showed that this fungal genus was present in a wide depth gradient, from 380 to 3474 m, indicating that it can be highly tolerant to gradients in temperature, dissolved oxygen, food supply, and the hydrostatic pressure associated with this change in depth. However, in six of the studied sediment cores *Metschikowia* was almost absent, pointing more to microhabitat variability.

The most abundant genus within Chytridiomycetes was *Rhizophidium* which can function as parasite and decomposer (Letcher et al., 2006; Kagami et al., 2007; Frenken et al., 2017), while the most abundant genera of Eurotiomycetes and Dothideomycetes were *Aspergillus* and *Cladosporium*, respectively. Within Agaricomycetes, the most abundant phylotype was related to the genus *Armillaria*. Previous studies have shown that *Aspergillus* and *Penicillium* are common inhabitants of deep-sea sediments; likewise, the presence of yeasts in this ecosystem has been frequently detected, but mainly related to genera such as *Pichia, Cryptococcus, Malassezia*, and *Rhodotorula* (Takishita et al., 2006; Zhang et al., 2015; Nagano et al., 2016, 2017; Grossart et al., 2019). To our knowledge, this is the first work showing a high abundance of *Metschikowia* in deep-sea ecosystems.

We used network analysis to explore possible relationships between eukaryotic microorganisms that coexist in deep marine sediments of Costa Rica (**Figure 4**). This technique allowed us to visualize positive associations not only between the members of the fungal taxa but also between fungi and other eukaryotes. For example, we confirmed the strong relationship between *Metschikowia*, the most abundant ASV from Saccharomycetes, and *Rhizophydium*, the most abundant ASV from Chytridiomycetes. Interestingly, in this yeast-dominated group, there were also associations with other eukaryotic ASVs belonging to Cercozoa, Ichthyosporea, Porifera, Annelida, and Cnidaria (**cluster 1, Figure 4**).

**Figure 4.**
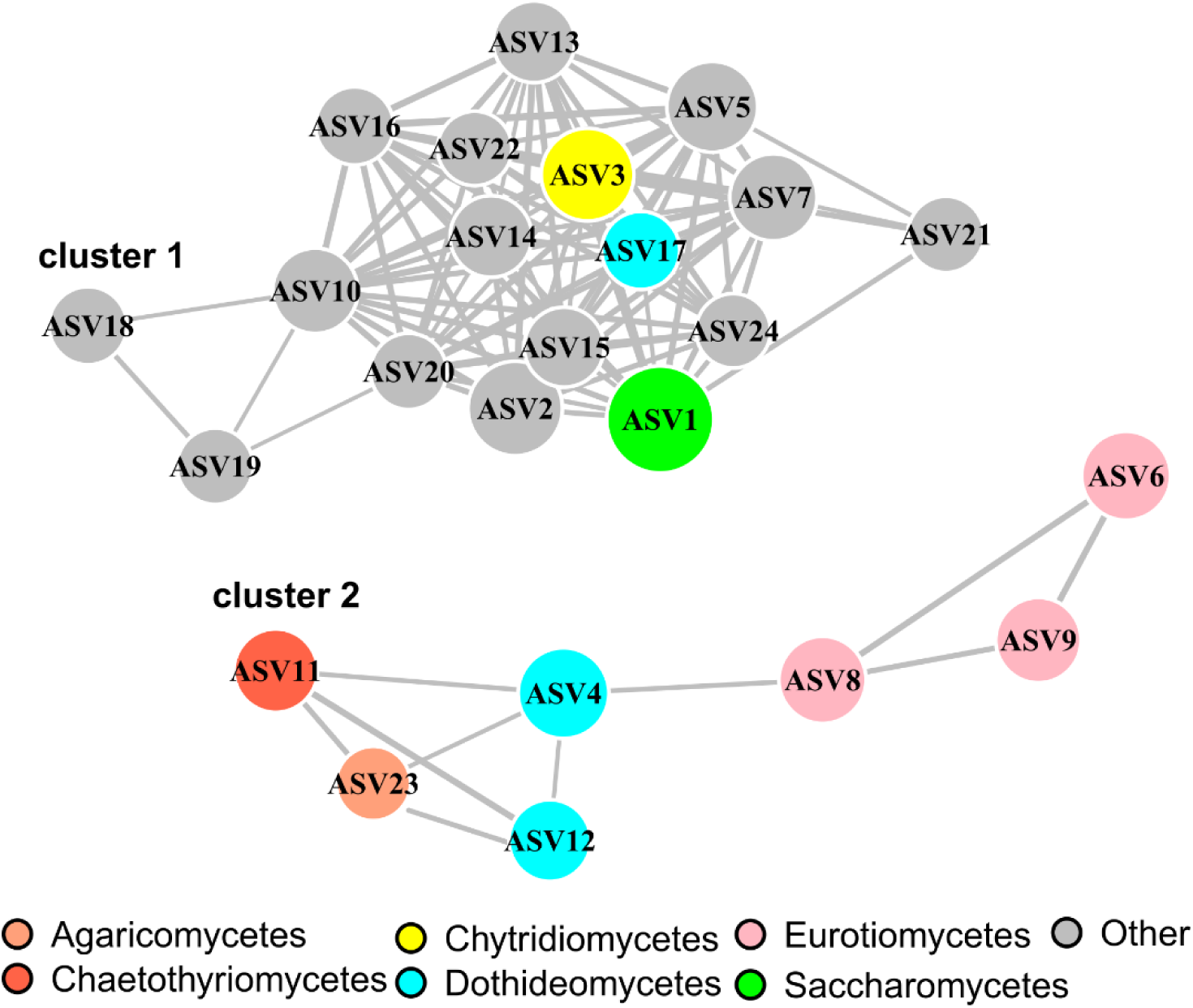
Network analysis highlighting the relationships between the groups of fungi and also with respect to other groups of eukaryotes present in 40 samples from deep-sea sediments. The analysis is based on the 24 most abudant eukaryotic ASVs (10 classified as fungi), which corresponded to nearly 70% of the total number of eukaryotic sequences. Colors of the nodes represent the taxonomic afiliation of the ASVs, while the size of the circles is proportional to their log-abundance. The network was generated and visualizaed with package igraph. The taxonomic classification of the ASVs at the genus level and kingdom level are shown as follows: ASV1: *Metschnikowia*(Fungi), ASV2: *Gymnophrys*(Cercozoa), ASV3: *Rhizophydium*(Fungi), ASV4: *Cladosporium*(Fungi), ASV5: *Pirum*(Ichthyosporea), ASV6: *Aspergillus*(Fungi), ASV7: *Pirum*(Ichthyosporea), ASV8: *Aspergillus*(Fungi), ASV9: *Aspergillus*(Fungi), ASV10: *Cryothecomonas*(Cercozoa), ASV11: *Exophiala*(Fungi), ASV12: *Neophaeosphaeria*(Fungi), ASV13: *Gymnophrys*(Cercozoa), ASV14: *Polymastia*(Porifera), ASV15: *Gymnophrys*(Cercozoa), ASV16: *Pirum*(Ichthyosporea), ASV17: *Acidomyces*(Fungi), ASV18: *Cossura*(Annelida), ASV19: *Tetrastemma*(Nemertea), ASV20: *Cryothecomonas*(Cercozoa), ASV21: *Pelagia*(Cnidaria), ASV22: *Cryothecomonas*(Cercozoa), ASV23: *Armillaria*(Fungi), ASV24: *Quadricilia*(Cercozoa).

In a second group, strong associations were detected between fungal genera such as *Cladosporium, Aspergillus, Exophiala*, and *Armillaria*, belonging to the classes Dothideomycetes, Eurotiomycetes, Chaetothyriomycetes, and Agaricomycetes, respectively. However, in this cluster, we did not detect associations with other eukaryotic ASVs that could point to co-occurrence with specific environmental settings (**cluster 2, Figure 4**).

The statistical analyzes, at the ASV level, showed no significant differences (Permanova, p> 0.05) in the structure of the communities according to variables depth, salinity, dissolved oxygen, pH, or between seep/non-seep areas nor between expeditions (**Supplementary Table 1, Supplementary Figure 2**). For example, we showed that depth (and, consequently, hydrostatic pressure) does not have an apparent effect on the composition of communities, given the wide distribution range of species. Furthermore, we observe that cluster 1 and cluster 2 inhabit sites whose temperature, salinity, dissolved oxygen and pH ranges overlap each other (**Table 2**). Therefore, it seems that the conditions of the deep waters are not limiting for the growth of the fungi and that there could be other variables influencing the composition of the communities whose effects should be further explored in future studies.

**Table 2.**
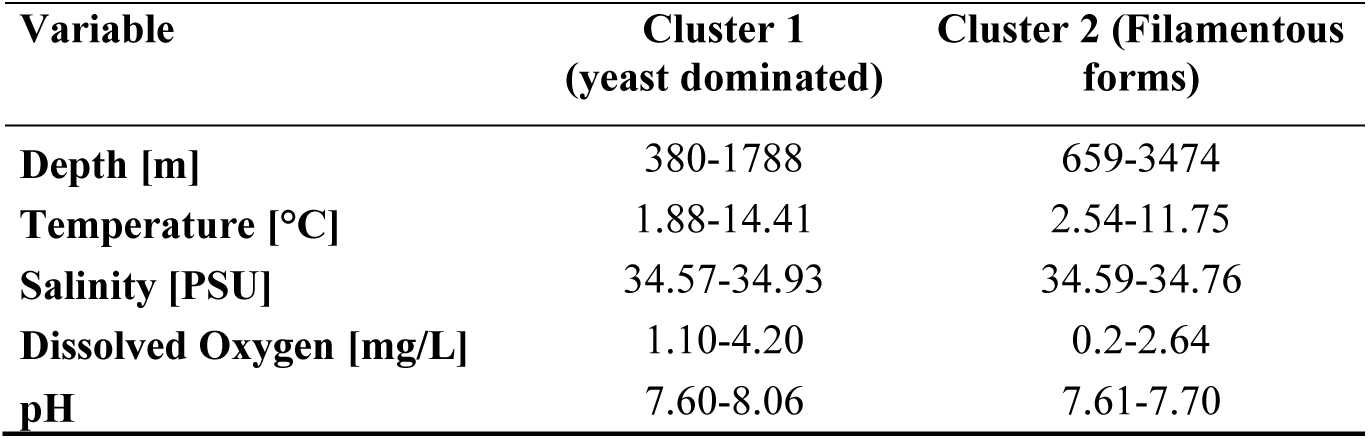
Range of values of the environmental variables for each of the fungal clusters identified

As an empirical observation note, samples that contained a higher proportion of mud were the ones that exhibited a higher abundance of Saccharomycetes (cluster 1). In contrast, sandy samples showed higher abundances of Eurotiomycetes and Dothideomycetes, which are filamentous fungi (cluster 2). This observation suggests a possible relationship between fungal morphology and its ability to colonize substrates of different textures. For example, yeasts may directly depend on the type and concentrations of organic matter found in the habitat, but could also perform fermentation processes in muddy sediments (Takishita et al., 2006; Kutty and Philip, 2008; Zhang et al., 2015; Taube et al., 2018).

We highlight the high prevalence of fungi in deep-sea sediments of the ETP of Costa Rica. The high abundance of yeasts like *Metschikowia* should be further studied using cultivation-dependent methods to provide better insights into the physiology, genomic makeup, and their contributions to global biogeochemical processes. Since it was difficult to distinguish the association of specific environmental variables with variations in the composition of fungal communities, particularly in the two clusters identified, further research will be necessary to determine how fungal communities in deep-sea waters are structured as well as to determine their ecological role in the largest biome on the planet.

## DATA AVAILABILITY

The dataset generated for this study can be found in NCBI Sequence Read Archive under accession PRJNA632873.

## AUTHOR CONTRIBUTIONS

KRJ, HPG, EEC and JC designed the study. JC and EEC collected the samples. KRJ, HPG, EEC and JC performed the analysis. KRJ wrote the manuscript. All authors helped to revise the manuscript.

## ACKNOWLEDGMENTS

We would like to thank all of the officers and crew of the RV *Atlantis* and RV *Falkor* for their assistance, and the pilots of the HOV *Alvin* and ROV *SuBastian* for their efforts in the collection of samples. We thank the Ministry of Environment and Energy of Costa Rica for granting the permits for the collection of samples. Sample processing was carried out with assistance from Odalisca Breedy and environmental data were compiled by Steve Auscavitch, Jay Lunden, and April Stabbins at Temple University. We also thank Lisa Levin and Greg Rouse at Scripps Oceanography, UC San Diego, La Jolla, CA, and Victoria Orphan at the California Institute of Technology, Pasadena, CA for their support during the development of this project. We acknowledge the support of the Deutsche Forschungsgemeinschaft and Open Access Publishing Fund of University of Potsdam

## SUPPLEMENTARY MATERIAL

**Supplementary Table 1.**
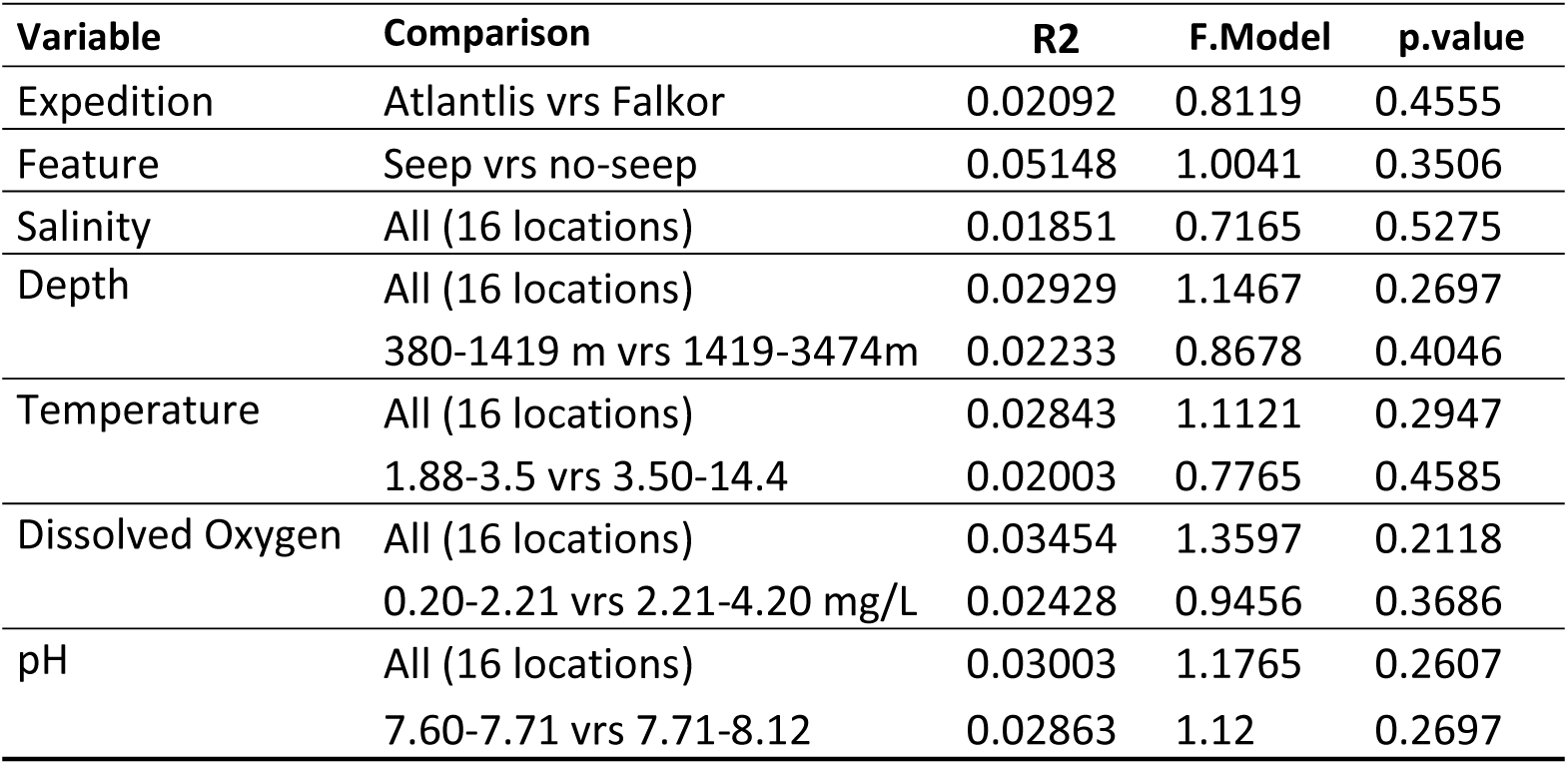
Statistical analysis of the fungal community composition related to different variables. The PERMANOVA tests were performed using function adonis2 and implemented in Vegan package. Data were normalized by converting the ASV counts into relative abundances. Binning of continuous variables Depth, Temperature, Dissolved Oxygen and pH was performed with package Hmisc.

| Variable | Comparison | R2 | F.Model | p.value |
| --- | --- | --- | --- | --- |
| Expedition | Atlantlis vrs Falkor | 0.02092 | 0.8119 | 0.4555 |
| Feature | Seep vrs no-seep | 0.05148 | 1.0041 | 0.3506 |
| Salinity | All (16 locations) | 0.01851 | 0.7165 | 0.5275 |
| Depth | All (16 locations) | 0.02929 | 1.1467 | 0.2697 |
|  | 380-1419 m vrs 1419-3474m | 0.02233 | 0.8678 | 0.4046 |
| Temperature | All (16 locations) | 0.02843 | 1.1121 | 0.2947 |
|  | 1.88-3.5 vrs 3.50-14.4 | 0.02003 | 0.7765 | 0.4585 |
| Dissolved Oxygen | All (16 locations) | 0.03454 | 1.3597 | 0.2118 |
|  | 0.20-2.21 vrs 2.21-4.20 mg/L | 0.02428 | 0.9456 | 0.3686 |
| pH | All (16 locations) | 0.03003 | 1.1765 | 0.2607 |
|  | 7.60-7.71 vrs 7.71-8.12 | 0.02863 | 1.12 | 0.2697 |

**Supplementary Figure 1.**
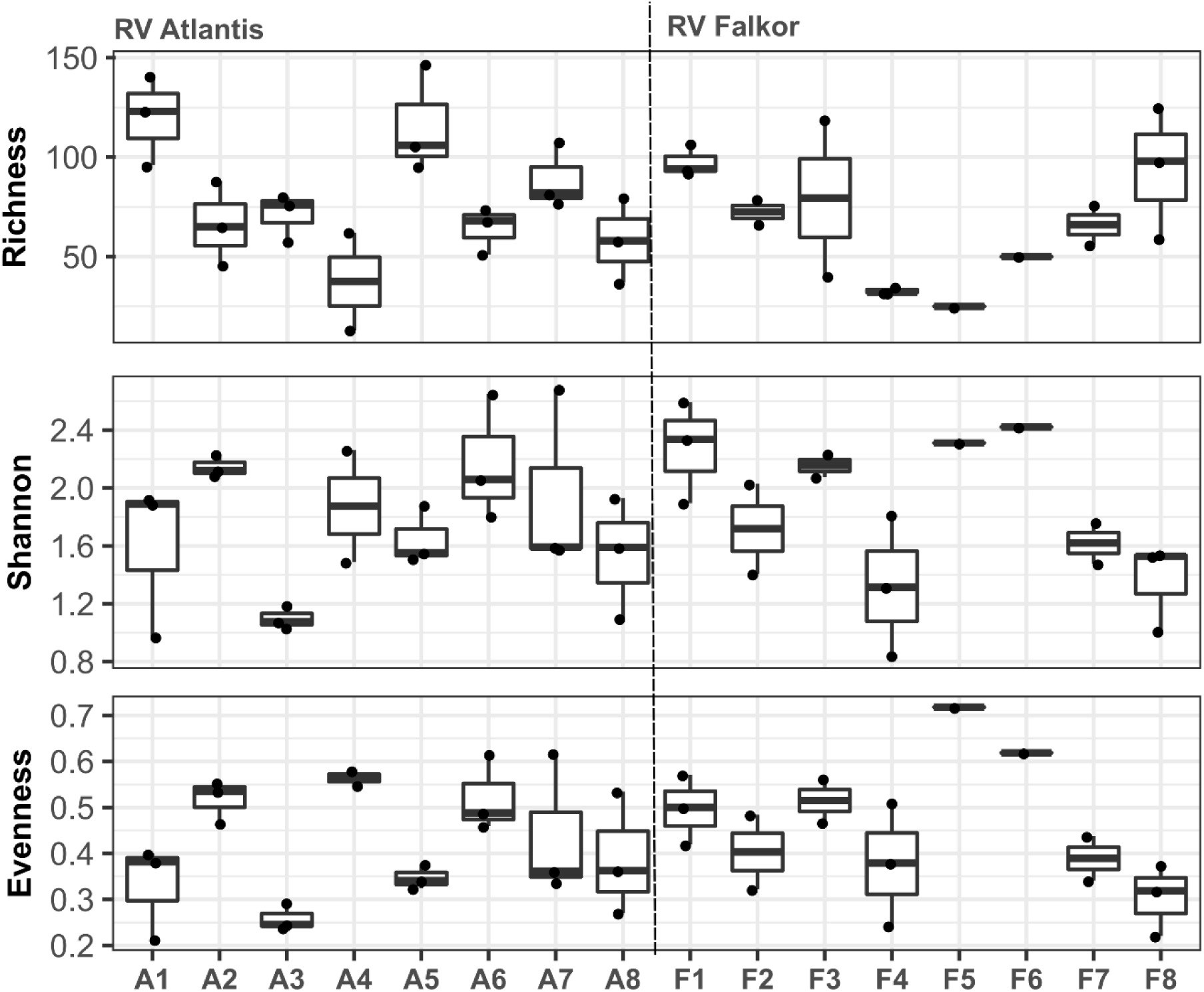
Boxplots of the alpha diversity estimations of the sampling points in deep-sea sediments of the Eastern Tropical Pacific.

**Supplementary Figure 2.**
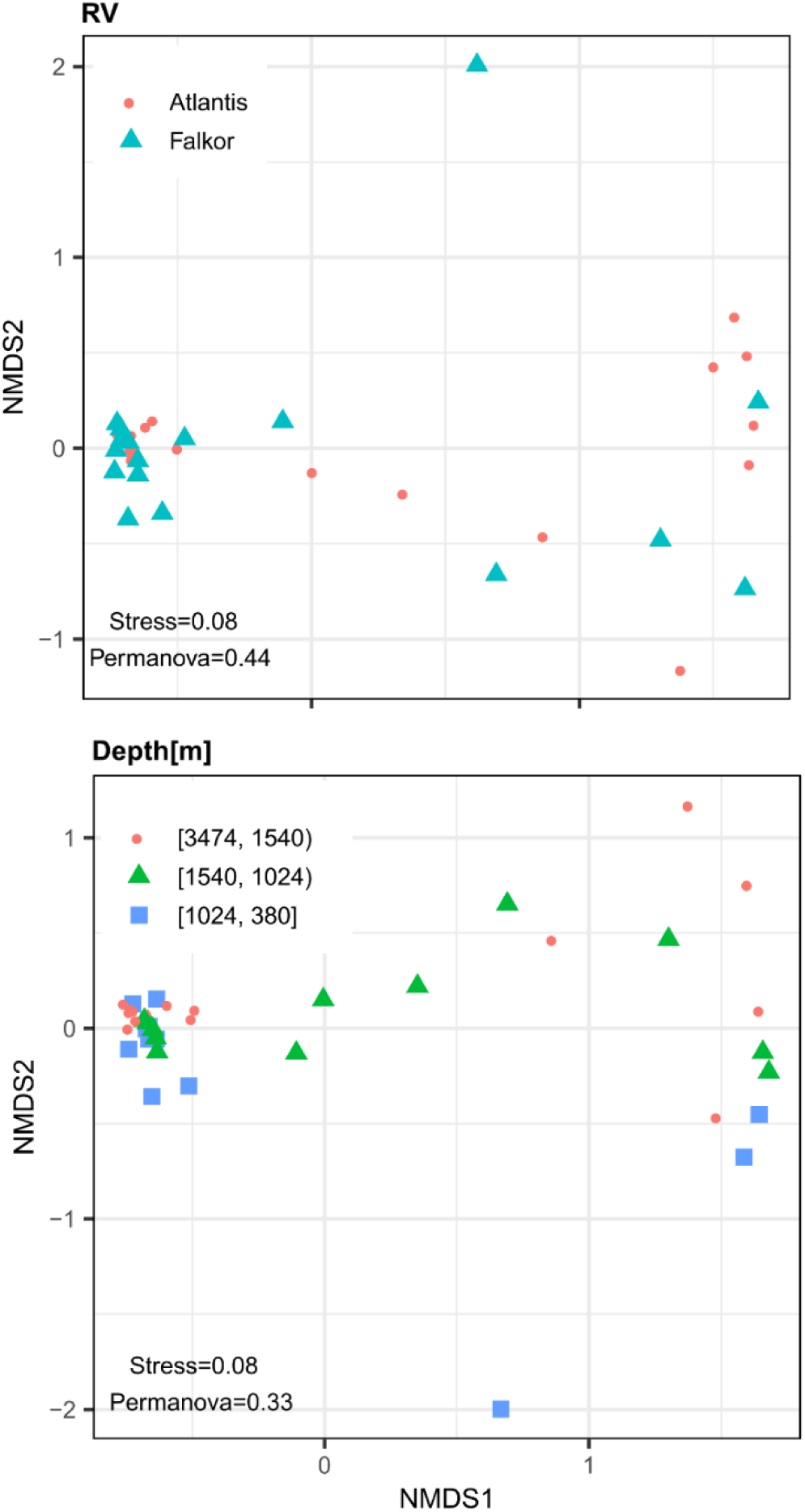
Non-metric multidimensional scaling (NMDS) analyses of the fungal communities in deep-sea sediments. The analysis include 40 samples from sediments obtained in a bathymetric gradient (from a depth of 380 to 3474 m) along two transects of about 1500 km length in the Eastern Tropical Pacific of Costa Rica. The upper panel shows the analysis by transect and the lower the analysis by depth. The stress values of the NMDS and the p value of the permanova analyses are also shown.

